# Human brain function during pattern separation follows hippocampal and neocortical connectivity gradients

**DOI:** 10.1101/2020.06.22.165290

**Authors:** Qiongling Li, Shahin Tavakol, Jessica Royer, Sara Larivière, Reinder Vos De Wael, Bo-yong Park, Casey Paquola, Debin Zeng, Benoit Caldairou, Danielle S. Bassett, Andrea Bernasconi, Neda Bernasconi, Birgit Frauscher, Jonathan Smallwood, Lorenzo Caciagli, Shuyu Li, Boris C. Bernhardt

**Author notes:** Corresponding Author: Boris C. Bernhardt, PhD, Multimodal Imaging and Connectome Analysis Lab, McConnell Brain Imaging Centre, Montreal Neurological Institute and Hospital, McGill University, Montreal, Quebec, Canada.

## Abstract

Episodic memory is our ability to remember past events accurately. Pattern separation, the process of of orthogonalizing similar aspects of external information into nonoverlapping representations, is one of its mechanisms. Converging evidence suggests a pivotal role of the hippocampus, in concert with neocortical areas, in this process. The current study aimed to identify principal dimensions of functional activation associated with pattern separation in hippocampal and neocortical areas, in both healthy individuals and patients with lesions to the hippocampus. Administering a pattern separation fMRI paradigm to a group of healthy adults, we detected task-related activation in bilateral hippocampal and distributed neocortical areas. Capitalizing on manifold learning techniques applied to parallel resting-state fMRI data, we could identify that hippocampal and neocortical activity patterns were efficiently captured by their principal gradients of intrinsic functional connectivity, which follows the hippocampal long axis and sensory-fugal cortical organization. Functional activation patterns and their alignment with these principal dimensions were altered in patients. Notably, inter-individual differences in the concordance between task-related activity and intrinsic functional gradients were correlated with pattern separation performance in both patients and controls. Our work outlines a parsimonious approach to capture the functional underpinnings of episodic memory processes at the systems level, and to decode functional reorganization in clinical populations.

## Introduction

Episodic memories are records of experiences and events in daily lives. Pattern separation is a core process in episodic memory, and is understood as the ability to orthogonalize similar patterns of external information into nonoverlapping representations (Bakker, Kirwan et al. 2008). Studying pattern separation, as well as the principles governing its localization in the brain, thus provides the opportunity to advance our understanding of the neural substrates of human episodic memory.

The medial temporal lobe, particularly the hippocampus, plays a key role in the formation and storage of episodic memories more generally (Milner, Squire et al. 1998) and in pattern separation specifically (Leal and Yassa 2018). Prior investigations of the hippocampus emphasized its role as a nexus that receives information from sensory and association cortices and combines this information with modulatory input from limbic and subcortical regions, leading to the formation of coherent representations of individual experiences, that are then projected back to several cortical regions (Lavenex and Amaral 2000). Task-based functional MRI (fMRI) studies have provided ample support for hippocampal activation during episodic memory tasks (Moscovitch, Cabeza et al. 2016). Patients with hippocampal lesions are known to present with difficulties in acquiring new episodic memories, and often forget memories of specific events (Rempel-Clower, Zola et al. 1996, Stefanacci, Buffalo et al. 2000). Previous task-based fMRI studies in humans compared brain activation during the presentation of novel or previously presented items against that for presentation of similar items, substantiating hippocampal activation during pattern separation (Bakker, Kirwan et al. 2008, Kyle, Stokes et al. 2015, Berron, Schutze et al. 2016).

Several, but not all of these studies (Reagh, Watabe et al. 2014, Pidgeon and Morcom 2016), suggested a role of specific hippocampal subdivisions in pattern separation, notably the dentate gyrus (DG) and specific Cornu Ammonis (CA) subfields, such as CA3 (Bakker, Kirwan et al. 2008, Berron, Schutze et al. 2016). Functional studies in both animals and humans suggest shifts in activity as a function of position on the hippocampal long axis, another dimension of hippocampal subregional specialization; in the context of pattern separation, however, this shift has not been extensively studied (Poppenk, Evensmoen et al. 2013, Strange, Witter et al. 2014, Collin, Milivojevic et al. 2015, Dalton, McCormick et al. 2019). Finally, despite much research focusing on the subregional specialization in the hippocampus, many studies emphasized that memory-related processes, like pattern separation, cannot be fully captured without reference to large-scale network effects. Particularly relevant studies implicate pattern separation-related functional activation in both higher order and sensory cortices (Pidgeon and Morcom 2016, Reagh, Murray et al. 2017). Collectively, findings suggest that pattern separation processes capitalize on subregional and network effects, motivating research interrogating both levels of neural organization.

Novel manifold analysis techniques applied to large-scale connectome datasets can identify principal axes of brain organization that describe smooth inter-regional transitions (henceforth, *gradients*). Applied to resting-state fMRI connectivity data, these techniques can capture shifts in connectivity patterns across both hippocampal subregions (Vos de Wael, Lariviere et al. 2018, Plachti, Eickhoff et al. 2019, Przezdzik, Faber et al. 2019) and neocortical areas (Margulies, Ghosh et al. 2016, Haak, Marquand et al. 2018). In the hippocampus, these techniques revealed two major axes of organization, with a primary gradient depicting the hippocampal long-axis and a secondary medio-lateral gradient that follows hippocampal infolding and subfield-to-subfield variations in microstructure (Vos de Wael, Lariviere et al. 2018, Plachti, Eickhoff et al. 2019). At the level of the neocortex, the principal functional gradient has repeatedly been shown to differentiate lower-level sensory systems from higher order association cortex (Margulies, Ghosh et al. 2016). As a main principle of functional cortical organization, gradients have also been leveraged to interrogate structure-function coupling (Huntenburg, Bazin et al. 2017, Paquola, Vos De Wael et al. 2019), brain development (Ball, Seidlitz et al. 2019, Lariviere, Vos de Wael et al. 2019, Paquola, Bethlehem et al. 2019), aging (Lowe, Paquola et al. 2019, Bethlehem, Paquola et al. 2020), and disease effects (Hong, Vos de Wael et al. 2019, Tian, Zalesky et al. 2019). Furthermore, gradients have proven useful in assessing distributed patterns of task-related neural activity underlying cognitive functions (Karapanagiotidis, Vidaurre et al. 2018, Murphy, Jefferies et al. 2018, Wang, Margulies et al. 2020). These findings suggest that gradient-based analysis may advance our understanding of principal organizational axes underlying neural function. In the context of episodic memory processes, stratification of functional activations along such hippocampal and neocortical gradients may provide a continuous and compact analytical space to interrogate subregional specialization as well as system-level integration in a unified framework.

A core goal of this analysis was to study pattern separation processes in adults during a task-based fMRI experiment, and to project activations into hippocampal and neocortical gradient spaces derived from parallel resting-state fMRI acquisitions. In addition to assessing this novel approach in healthy adults, we also investigated a cohort of well-defined patients with mesial temporal lobe epilepsy who present with variable degrees of hippocampal structural anomalies and behavioral deficits in pattern separation. As such, we could capitalize on this condition as a human “lesion model” for functional impairment and associated functional network reorganization following mesiotemporal damage.

## Results

We studied 26 healthy adults (15 males; 21-44 years), recruited by advertisement, as well as 14 patients with drug-resistant mesial left temporal lobe epilepsy (6 males; 19-56 years) based on a multimodal neuroimaging paradigm that involved high-resolution structural MRI, resting-state functional MRI, as well as task-based assessments of pattern separation (for details, see materials and methods). Studying healthy individuals, behavioral pattern separation (BPS) scores were calculated as the pattern separation rate ***P***(***similar***|***lure***) corrected for similar bias rate ***P***(***similar***|***novel***) based on the responses given in the second phase of the fMRI task. BPS scores followed an approximate normal distribution with mean=0.39 and standard error=0.20 in our healthy controls (**Figure 1A**). Voxel-wised general linear models mapped pattern separation activation based on the contrast (***similar***|***lure***) vs (***similar***|***novel***) as in prior work (Stark, Yassa et al. 2013). We observed main effects seen in bilateral hippocampus as well as distributed cingulate, frontal, temporal, and parietal areas (*p*<0.001, uncorrected, cluster size >5; **Figure 1B**).

**Figure 1.**
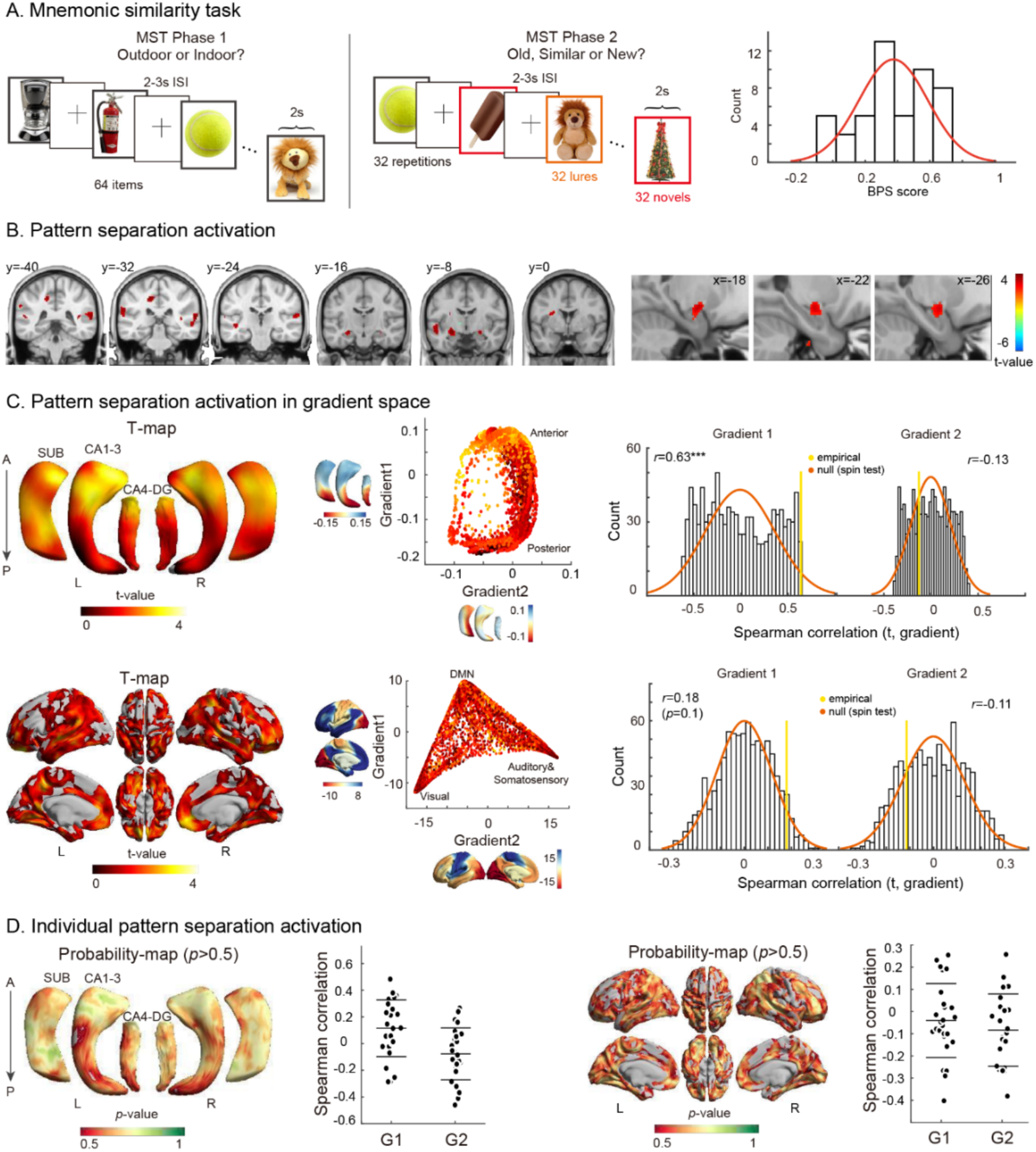
Mapping pattern separation activation to connectome gradient space. **A)** The mnemonic similarity task (MST) design and behavioral pattern separation (BPS) responses in healthy controls. **B)** Voxel-wise pattern separation mapping in controls. Findings are shown at p<0.001 uncorrected. **C)** Gradient analysis. Pattern separation activations are mapped to neocortical and hippocampal subfield surfaces (*left*) and projected to functional gradient space (*center*). The first neocortical gradient depicts a transition from unimodal to transmodal areas and the second gradient differentiates visual from somatomotor cortices (Margulies, Ghosh et al. 2016). The first hippocampal gradient follows its long axis and the second its infolding (Vos de Wael, Lariviere et al. 2018). The correlation between gradient maps and activation maps can be calculated, and statistical significance can be determined using nonparametric spin tests that account for spatial autocorrelation (Alexander-Bloch, Shou et al. 2018). **D)** The analysis was repeated at a single subject level, assessing correlations between individualized activation patterns and functional gradients.

We then projected the task-based fMRI activation patterns to low dimensional manifold spaces spanned by intrinsic connectivity gradients in neocortical and hippocampal subregions (**Figure 1C**). Gradients were derived from parallel resting-state fMRI acquisitions in the same individuals, following established methods (Vos de Wael, Benkarim et al. 2020). Gradients were consistent with prior work on the human connectome project dataset (Margulies, Ghosh et al. 2016, Vos de Wael, Lariviere et al. 2018): In the hippocampus, the first and second gradient described an antero-posterior and medio-lateral differentiation (Vos de Wael, Lariviere et al. 2018). In the neocortex, the first and second gradient described a unimodal-transmodal and visual-somatosensory transition, respectively (Margulies, Ghosh et al. 2016)

In the hippocampus, an assessment of the correlation between task based activation patterns and functional gradients revealed a strong and significant correspondence between activations and functional gradients. Specifically, activation strength correlated strongly with the first principal gradient that differentiated anterior-posterior hippocampal divisions, with stronger activation in anterior segments (*r* = 0.63; *spin* − *test, p* < 0.001). In the neocortex, we observed a marginal positive correlation between activation strength and the first unimodal-to-transmodal gradient, with significances determined using non-parametric spin tests that control for spatial autocorrelations (*r* = 0.18; *spin* − *test, p* = 0.1). These findings indicate that activations tended to be stronger at the transmodal apex compared to the unimodal end.

We also mapped individual subject activations (*i.e*., positive t-values obtained from the first level analysis) into gradient space and repeated the above analyses. Findings were consistent with the group level findings, indicating positive activations in the majority of participants in similar regions as those identified by the group analysis. Furthermore, we obtained a consistently positive correlations between subject-specific activations and the first hippocampal gradient (*t*(26) = 2.75, *p* = 0.01).

### Functional reorganization in patients with hippocampal anomalies

Studying a cohort of patients with temporal lobe epilepsy, we investigated functional reorganization in individuals with variable degrees of hippocampal structural pathology and behavioral deficits in pattern separation. Although patients and controls did not significantly differ by age or sex, the former showed reduced hippocampal volumes in the left hemisphere compared to controls, as well as increased inter-hemispheric volumetric asymmetry (*t*(36) = −3.91, *p* < 0.001, **Supplementary Figure S1**).

When comparing behavioral performance between patients and controls, we detected reduced pattern separation abilities in patients. Whereas controls successfully labeled 51.8±16% of lure trials as “similar”, patients did so on for only 33.3±16% trials (*t*(36) = 2.72, *p* = 0.02, FDR-corrected, **Figure 2A**). Critically, performance on other trial types did not differ across groups, and patients also showed intact pattern completion, *i.e*., labeling lure trials as “old” (see **Supplementary Table S1**). When the behavioral pattern separation score was calculated as *BPS* = *p*(*similar*|*lure*) − *p*(*similar*|*novel*) *i.e*., when scores were adjusted for the similar bias rate, patients still showed significant impairment compared to controls (*t*(36) = 2.73, *p* = 0.02, FDR-corrected). Findings were virtually identical after repeating between-group contrasts after controlling for age and sex.

**Figure 2.**
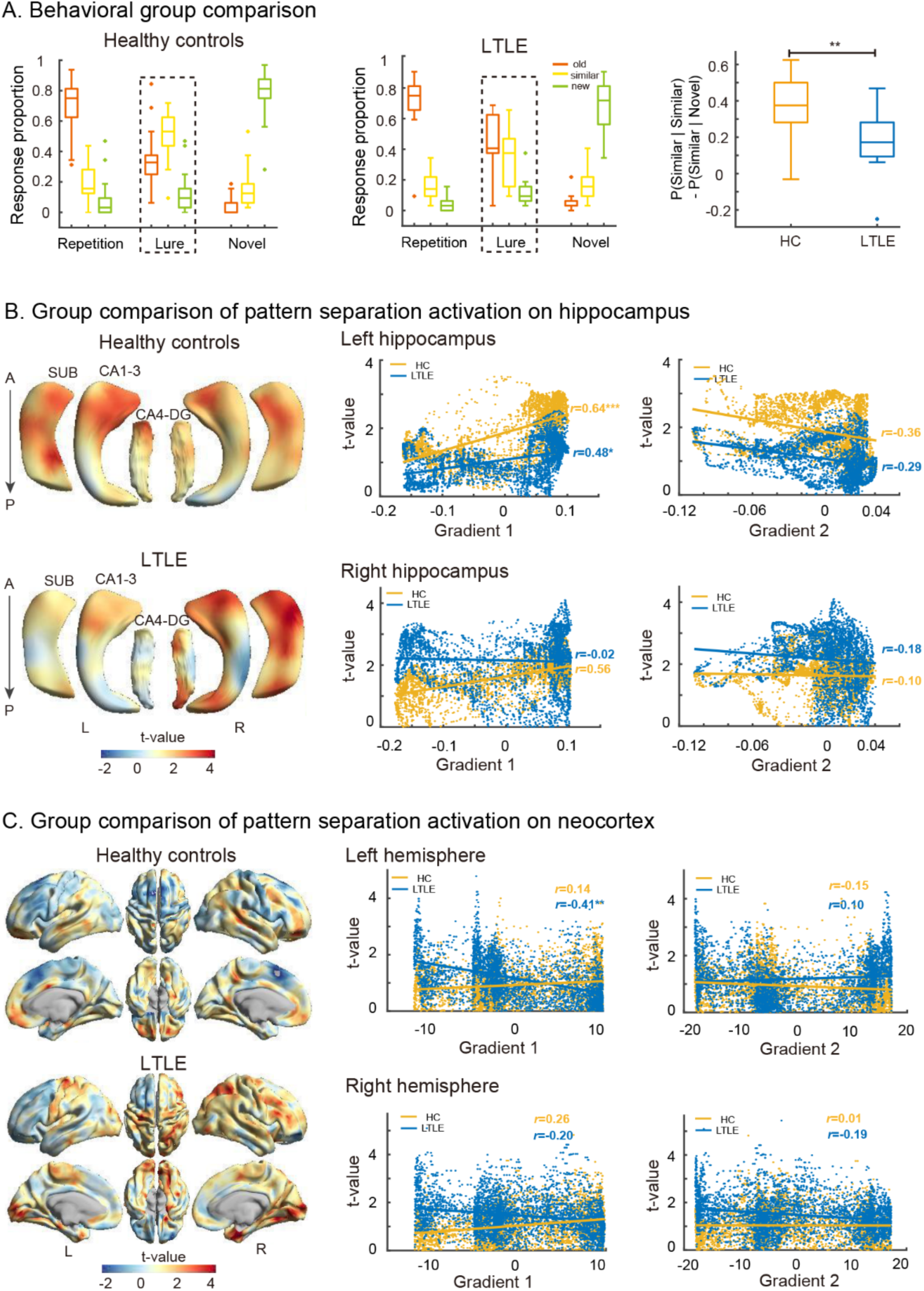
Perturbations in structure-function relations in neurological patients with hippocampal damage. **A)** Response proportions and BPS score in both patients and controls and group comparison controlled for age and sex, showing reduced pattern separation in patients. **B) & C)** Correlations between pattern separation activations with both groups and connectome gradients, showing that the topography of functional activations in patients did not follow connectome gradients the same way as in controls, specifically with respect to the first gradient in hippocampal and neocortical regions.

Connectome gradient stratification was also useful in visualizing between-group differences in how functional activations followed hippocampal and neocortical topographies (**Figure 2B-C**). Compared to controls, patients presented with different activation-gradient relationships, specifically with respect to the first gradient in both hippocampal and neocortical areas. Correlations between functional activation patterns and the second gradient were similar in both patients and controls, despite overall lower activations in patients in the ipsilateral hippocampus and stronger activations contralaterally (**Figure 2B-C**).

Formally, while activation of the left hippocampus (*i.e*., ipsilateral to the seizure focus) in patients was also positively correlated with the first gradient (*r* = 0.48, *spin* − *tests p* < 0.05), correlations were weaker than in controls (*r* = 0.64, *spin* − *tests p* < 0.001). This finding was also confirmed using a direct comparison of correlation coefficient magnitudes. This attenuation on the side of pathology co-occurred with possible reorganization effects within the right (*i.e*., contralateral) hippocampus. Here, while controls showed a similar positive alignment between functional activations and the first gradient (*r* = 0.56, *spin* − *tests p* < 0.05), patterns disappeared in patients (*r* = −0.02, *spin* − *tests p* > 0.05), who presented stronger activations in both ends (left: *p* > 0.05, right: *p* > 0.05).

In the neocortex, patients showed an opposite correlation between activations and the principal gradient (left: *r* = −0.41, *spin* − *tests p* < 0.01; right: *r* = −0.20, *p* = 0.1), compared to healthy controls (left: *r* = 0.14, *spin* − *tests p* = 0.1; right: *r* = 0.26, *spin* − *tests p* = 0.1). Unlike controls, patients showed stronger activation at the unimodal relative to the transmodal end in both left and right hemispheres (left: *p* = 0.001; right: *p* = 0.037).

### Behavioral associations

To explore behavioral associations, we weighted subject specific *t*-values from the task-based fMRI experiment relative to the neocortical and hippocampal principal gradient (**Figure 3)**. This procedure provided a personalized loading scores, with high loadings indicating that activity patterns followed the gradients, and low loadings indicating the opposite. Subject-specific loadings were used as regressors for inter-individual differences in behavioral pattern separation score, controlling for age and sex across both patients and controls. While left hippocampal loadings were correlated to BPS scores (*r* = 0.40, *p* = 0.01), right loadings were not (*r* = 0.24, *p* = 0.13). Considering neocortical loadings, marginal associations were observed in both the left and right hemispheres (left: *r* = 0.24, *p* = 0.13; right: *r* = 0.29, *p* = 0.07)

**Figure 3.**
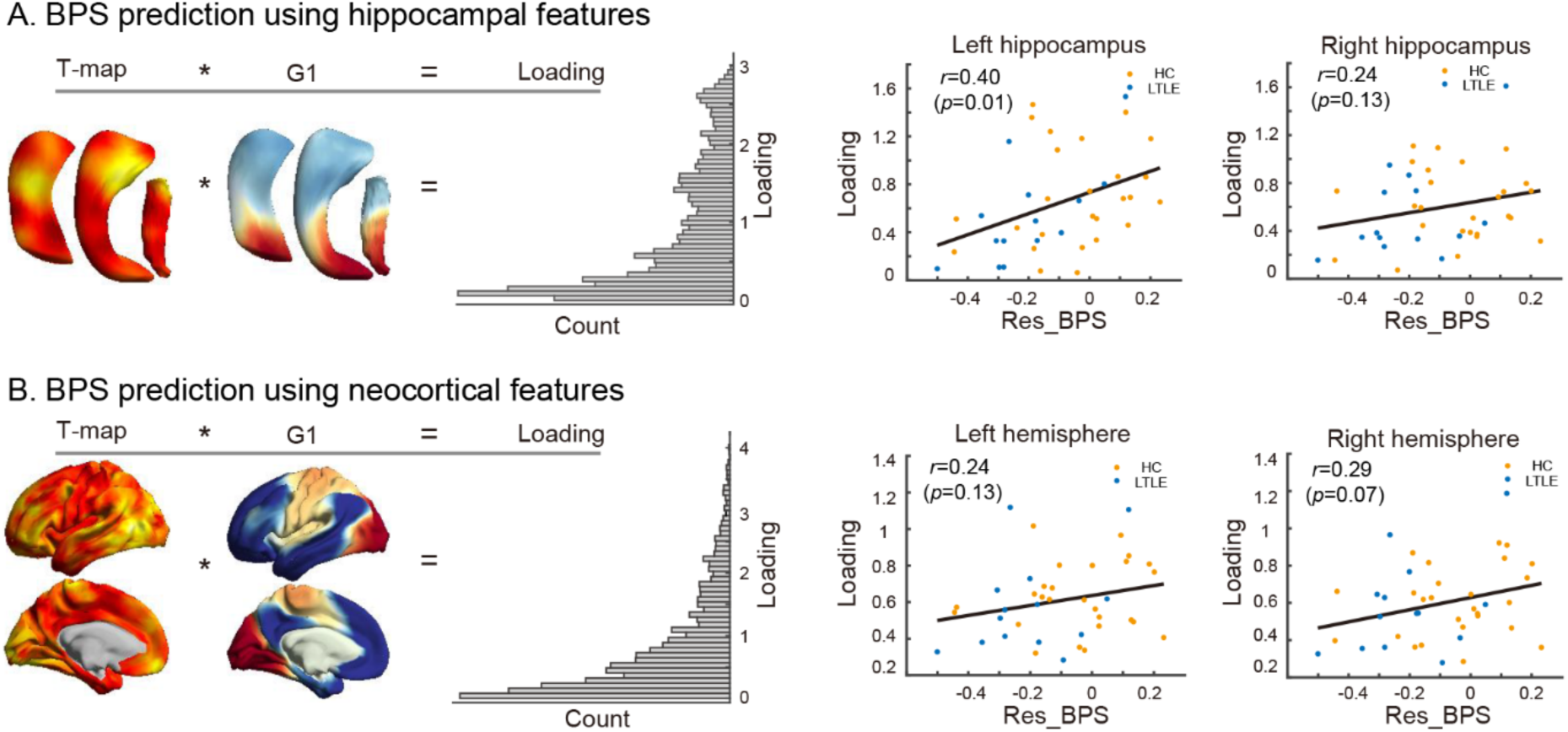
Behavioral associations. **A)** A subject specific loading score was computed by weighting individual t-statistical maps from the pattern separation activation by normative gradient maps in the hippocampus (*left*). We then assessed the correlation between these loadings and BPS scores, after controlling for age and sex for both hippocampal and neocortical regions *(right)*. **B)** Equivalent analysis for the neocortex.

## Discussion

Memory processes are known to critically engage the hippocampus in concert with distributed brain networks. Here, we capitalized on connectome manifold learning techniques to identify intrinsic functional gradients, and to identify compact signatures of hippocampal and neocortical functional activations during pattern separation. Studying a cohort of healthy young adults using multimodal imaging, we observed that task-based activations during a pattern separation paradigm were strongly captured by intrinsic functional connectivity gradients in the hippocampus, which recapitulates its long axis (Vos de Wael, Lariviere et al. 2018, Plachti, Eickhoff et al. 2019, Przezdzik, Faber et al. 2019), and marginally in the neocortex, where functional activity tended to increase towards the transmodal apex (Margulies, Ghosh et al. 2016). Translating our approach to patients with drug-resistant temporal lobe epilepsy and mesiotemporal lesions, we could furthermore assess the functional impact exerted by structural anomalies. Specifically, we identified distorted relations between connectome gradients and functional activations in hippocampal as well as neocortical networks, relative to controls. Finally, our approach delivered a personalized fMRI signature score indexing the alignment of functional activations to connectome gradients. This metric was found to be associated with behavioral performance in both patients and controls, and particularly effective in the hippocampus. Collectively, these findings identify topographic underpinnings episodic memory processing in humans, and chart potential mechanisms governing functional reorganization in patients with a neurological disease who present with damage to pivotal nodes in the human memory network.

Recent years have seen an increase in the application of manifold learning techniques to capture smooth inter-regional transitions in brain function, microstructure, and connectivity (Burt, Demirtaş et al. 2018, Haak, Marquand et al. 2018, Huntenburg, Bazin et al. 2018, Fulcher, Murray et al. 2019, Paquola, Vos De Wael et al. 2019, Shine, Breakspear et al. 2019, Vos de Wael, Benkarim et al. 2020). Leveraging resting-state fMRI, these techniques have visualized connectivity gradients in both hippocampal subregions (Vos de Wael, Lariviere et al. 2018, Plachti, Eickhoff et al. 2019, Przezdzik, Faber et al. 2019) as well as neocortical areas (Margulies, Ghosh et al. 2016, Haak, Marquand et al. 2018). In contrast to traditional whole-brain voxel-wise activation analysis (Pidgeon and Morcom 2016) or region of interest approaches that focus on the whole hippocampus or its subfields (Berron, Schutze et al. 2016), gradient mapping techniques capture salient organizational axes of intrinsic function in a low-dimensional space. This estimate is potentially less affected by subtle inter-individual variations and challenges in reliably defining specific subregions, such as hippocampal subfields themselves (Yushkevich, Wang et al. 2010, Kulaga-Yoskovitz, Bernhardt et al. 2015). A growing number of studies have capitalized on gradient stratification to assess changes in functional activation patterns following a specific task (Murphy, Jefferies et al. 2018, Karapanagiotidis, Vidaurre et al. 2019), to visualize spatial trends of neurodegenerative deposits and cortical thinning in aging (Lowe, Paquola et al. 2019, Bethlehem, Paquola et al. 2020), and to assess molecular underpinnings of neuroimaging findings (Burt, Demirtaş et al. 2018, Vogel, La Joie et al. 2020).

Here, we projected task-based functional activations into this alternative space governed by connectivity gradients in both neocortical and hippocampal regions. An adapted version of the MST paradigm was used, as it allows for sensitive functional mapping and the calculation of behavioral pattern separation scores at the group and single subject levels (Stark, Yassa et al. 2013, Stark, Kirwan et al. 2019). Prior task-based fMRI analyses identified both hippocampal and isocortical activations during pattern separation (Bakker, Kirwan et al. 2008, Lacy, Yassa et al. 2011, Berron, Schutze et al. 2016, Pidgeon and Morcom 2016, Reagh, Murray et al. 2017); notably, these studies generally carried out conventional region-of-interests analyses, either at the level of the entire hippocampus (Bakker, Kirwan et al. 2008, Pidgeon and Morcom 2016) or hippocampal subfields (Bakker, Kirwan et al. 2008, Lacy, Yassa et al. 2011, Reagh, Watabe et al. 2014), or ran unconstrained voxel-wise analysis (Lacy, Yassa et al. 2011, Reagh, Murray et al. 2017). In our work, gradient stratification of activations during pattern separation showed strong alignment with the principal hippocampal gradient, and a marginal alignment with the neocortical gradient, anchoring our findings in established models of hippocampal long axis specialization (Poppenk, Evensmoen et al. 2013, Strange, Witter et al. 2014, Vos de Wael, Lariviere et al. 2018, Dalton, McCormick et al. 2019, Plachti, Eickhoff et al. 2019, Przezdzik, Faber et al. 2019) and cortical hierarchies (Mesulam 1998, Chaudhuri, Knoblauch et al. 2015, Chanes and Barrett 2016). Overall, our findings provide a compact model to conceptualize functional participation of mesiotemporal and neocortical subregions in pattern separation.

Our findings suggest increased functional activity towards to the anterior hippocampal segments and densely connected transmodal association cortices in cognitive processes related to pattern separation. In the hippocampus, prior work on pattern separation has already investigated subregional specialization with respect to hippocampal subfields, particularly CA3/DG. To the authors knowledge, these processes have however not been extensively studied relative to the hippocampal long axis, which is considered as another important dimension of hippocampal subregional organization (Poppenk, Evensmoen et al. 2013, Vos de Wael, Lariviere et al. 2018, Dalton, McCormick et al. 2019). Our results seem to echo findings from other memory tasks, that suggest a distribution of neural processing across different hippocampal segments and different levels of the cortical hierarchy. Indeed, a prior study using a scene repetition task observed distinct familiarity and repetition-related recognition signals in the DG/CA3 (Reagh, Watabe et al. 2014), which were dissociated along the hippocampal longitudinal axis. Further work associated gist memory with increased anterior hippocampal activity (Gutchess and Schacter 2012). An established model of hippocampal long axis specialization (Poppenk, Evensmoen et al. 2013) posits that hippocampal representations gradually vary along its long-axis, with broader anterior representations and sharper posterior representations; this differentiation may be relevant in the context of the pattern separation, where similar items need to be adequately discriminated from old items. Hippocampal subregions are likely to carry out these computations in concert with other brain networks (Pidgeon and Morcom 2016, Reagh, Murray et al. 2017). Consistent with these prior studies, we also observed neocortical participation in pattern separation. These were localized in both higher-order (e.g., default mode) apex regions in prefrontal and midline cortices, but also inferior temporal regions and occipital-temporal and that participate in ventral visual streams. The interplay between hippocampal activation shifts along the long axis, and recruitment of sensory as well as integrative cortical areas may reflect the contribution of these networks in visual processing (Ettlinger 1990, Goodale and Milner 1992) together with higher order transmodal cortices involved in self-related cognition (Northoff 2011), mental time travel (Schacter, Addis et al. 2012, Karapanagiotidis, Bernhardt et al. 2017), as well as cognitive control (Badre and Wagner 2007). These macroscale findings could potentially reflect computational accounts justified by evidence in animals (Rolls 2016), whereby hippocampal nodes invoke a reverse hierarchical series of cortical pattern association networks, implemented through hippocampal-cortical connections, to perform a form of pattern generalization, and to retrieve complete patterns stored in higher order areas.

Several experimental studies in non-human animals (McTighe, Mar et al. 2009) and behavioral assessments in neurological patients (Yassa, Stark et al. 2010, Fujii, Saito et al. 2014, Leal, Tighe et al. 2014, Semenova 2015, Reyes, Holden et al. 2018) have demonstrated deficits in pattern separation following hippocampal lesions. Here, we assessed patients with left mesial temporal lobe epilepsy who present with variable degrees of hippocampal pathology, using an identical neuroimaging and analysis paradigm as in our healthy controls. At the behavioral level, our young and middle-aged patients had a lower pattern separation score, giving more “old” responses and fewer “similar” responses to lure items than healthy controls, while showing unimpaired performance on novel and repeated items. This behavioral deficit is generally consistent with mnemonic findings in the epilepsy neurocognitive literature. For example, temporal lobe epilepsy patients typically do not perform as well as healthy individuals on visual and verbal memory tasks (Hermann, Seidenberg et al. 1997, Tramoni-Negre, Lambert et al. 2017), and recent work has also demonstrated impaired spatial pattern separation performance (Reyes, Holden et al. 2018). At the level of brain activations, we observed decreased ipsilateral functional activations, together with activity increases in the contralateral hemisphere. The involvement of other cortical regions and contralateral hippocampal regions in patients with left side hippocampal anomalies is consistent with prior findings (Bonnici, Sidhu et al. 2013, Sidhu, Stretton et al. 2013), and may overall indicate a broader extent of memory networks in patients with damage of core nodes in episodic memory circuits, indicative of functional reorganization associated to suboptimal memory performance. Interestingly, the pattern separation deficit in the young and middle aged epileptic patients mirrored findings in healthy adults aged greater than 70 years (Yassa, Lacy et al. 2011, Doxey and Kirwan 2015), a finding potentially in line with accounts suggesting that epileptic patients may experience accelerated brain aging (Bell, Lin et al. 2011, Bernhardt, Kim et al. 2013, Caciagli, Bernasconi et al. 2017, Galovic, Baudracco et al. 2019). A neurobiological framework was proposed to explain aging related impairments in pattern separation (Wilson, Gallagher et al. 2006), where reduced processing within the hippocampal circuitry, coupled with a decrease in cholinergic input, results in reduced pattern separation performance. Interestingly, low-performing patients in the current study activated their right hippocampus to a greater extent than healthy controls, consistent with studies of healthy elderly people (Miller, Celone et al. 2008, Yassa, Lacy et al. 2011).

As a limiting factor, we note that our sample size was modest and that we analyzed 3T imaging data only, which was however based on advanced and accelerated structural and functional sequences that can be readily integrated into studies in both healthy and diseased populations. Moving towards higher field strengths, such as 7T or beyond, may lead to improvements in spatial resolution and more fine grained functional mapping across different subregions (Berron, Schutze et al. 2016). One the other hand, successful scanning at these platforms will need to mitigate increases in susceptibility artifacts, that can considerably affect temporal lobe image fidelity. As our results furthermore show, our 3T framework already allows for robust and anatomically grounded stratification of functional activations across major axes of hippocampal and neocortical activation, and can anchor these findings relative to established dimensions of brain function and hierarchical processing (Huntenburg, Bazin et al. 2018). In addition to group-level analyses, the gradient wise coordinate system allowed us to explore across-subject consistency of functional activations, confirming a main role of the hippocampal long axis in pattern separation specifically. We furthermore calculated a subject-specific gradient *loading* score, an index of alignment of subject-specific functional activations with principal gradients. Notably, this approach revealed that individual behavioral separation ability was positively correlated with both left hippocampal loadings, and marginally with bilateral neocortical loading scores, in a sample composed of both patients with epilepsy and controls. These findings further support that stratification of brain functional activity from connectivity-based dimensions may provide a principled and compact way to obtain brain measures that are behaviorally and clinically relevant.

## Materials and Methods

### Participants

We studied 26 healthy adults (15 males; 21-44 years; mean±SD age=31.1±7.31 years, 1/25 left-/right-handed), recruited by advertisement, as well as 14 patients with drug-resistant left temporal lobe epilepsy (6 males; 19-56 years; mean±SD years=35.9±14.66, 2/12 left-/right-handed). Demographic and clinical data were obtained through interviews with patients and their relatives. Epilepsy diagnosis and lateralization of the seizure focus were determined by a comprehensive evaluation by the clinical team based on detailed history, neurological examination, review of medical records, prolonged video-EEG recordings, clinical MRI, neuropsychology assessment in all patients, and FDG-PET in a patient subgroup. There were no patients with mass brain lesions (malformations of cortical development, tumor, or vascular malformations), nor with history of traumatic brain injury or encephalitis. All patients had a left sided seizure focus. Our protocol received approval from the Research Ethics Board of the Montreal Neurological Institute and Hospital. All participants provided their informed consent in writing. See **Supplementary Table S1** for socio-demographic and clinical information on the patient and control cohorts included in this study.

### Mnemonic Similarity Task

The mnemonic similarity task (MST) was adapted from a previous study (Stark, Yassa et al. 2013), with stimuli taken from the original experiment (https://faculty.sites.uci.edu/starklab/mnemonic-similarity-task-mst/) and administered inside the scanner. It consisted of two phases (**Figure 1A**). In phase 1, participants were shown 64 color photographs of everyday objects on a white background and they gave an indoor-vs-outdoor judgment for each picture (lasting 2 seconds) via a button press. In phase 2, which occurred about 7 minutes after the encoding task, participants engaged in a recognition memory test, during which they identified each item as “old”, “similar” or “new” via a button press. Trials in phase 2 consisted of 32 novel items that were not seen before (novel), 32 similar items that were similar to those seen in phase 1 (lure), and 32 old items that were exactly the same in phase 1 (repetition). There was a fixation period lasting 2-3 seconds between each consecutive pair of items, resulting in a jittered inter-stimulus interval (ISI). The behavioral pattern separation (BPS) score was computed as the pattern separation rate *p*(*similar*|*lure*) corrected for similar bias rate *p*(*similar*|*novel*), thus resulting in *BPS* = *p*(*similar*|*lure*) − *p*(*similar*|*novel*).

### MRI acquisition

MR images were acquired using accelerated sequences on a Siemens Magnetom 3T PrismaFit scanner with a 64-channel head coil. Two T1-weighted images with identical parameters were acquired using a 3D MPRAGE sequence (TR=2300ms, TE=3.14ms, TI=900ms, flip angle=9°, FOV=256 × 256, matrix size=320 × 320, leading to a resolution of 0.8 × 0.8 × 0.8 *mm*^3^; ipat = 2; with an acquisition time of 6 minutes and 44 seconds). Resting-state fMRI data were acquired using a 2D echo planar imaging (EPI) sequence (TR=600ms, TE=30ms, flip angle=50°, FOV = 240 × 240, matrix size = 80 × 80, leading to a resolution of 3 × 3 × 3 *mm*^3^; multiband acceleration factor = 6; with an acquisition time of 7 minutes and 6 seconds). Participants were instructed to keep their eyes open, fixate on a cross centrally presented on a white screen, and not fall asleep. Task fMRI used a 2D EPI sequence with parameters identical to the resting-state fMRI (acquisition time of 5 minutes and 37 seconds for phase 1, 8 minutes and approximately 30 seconds for phase 2).

### MRI preprocessing

#### a) Structural MRI processing

Images underwent intensity non-uniformity correction and the two repeated scans were co-registered and averaged. Following brain extraction and segmentation of subcortical structures, preprocessed images were nonlinearly registered to MNI152 space and cortical surfaces were extracted using recon-all in FreeSurfer 6.0 (Dale, Fischl et al. 1999, Fischl, Sereno et al. 1999, Fischl, Sereno et al. 1999). To improve interindividual correspondence, cortical surfaces were aligned to the hemisphere-symmetric Conte69 template (Van Essen, Glasser et al. 2011). The hippocampus was automatically segmented into CA1-3, CA4-DG, and subiculum using our previously published and validated surface patch-based algorithm (Bernhardt, Bernasconi et al. 2016, Caldairou, Bernhardt et al. 2016). Hippocampal subfield surfaces were parameterized using a spherical harmonics approach (SPHARM-PDM). A Hamilton-Jacobi approach was used to create a medial sheet, representing the core of the subfield with a minimal partial volume effect. Spherical harmonics parameters on the outer hull were propagated to their corresponding medial sheet locations along a Laplacian field, thereby improving across subject correspondence.

#### b) Task-based fMRI

We used SPM12 (Penny, Friston et al. 2011) (version 6470, https://www.fil.ion.ucl.ac.uk/spm/software/spm12/) for MATLAB for all task-based fMRI processing. We first generated field maps from the AP-PA blip pairs and applied geometric distortion correction using the FieldMap toolbox (Hutton, Bork et al. 2002). Unwarped images were aligned to the first image using a rigid body transformation, followed by a linear co-registration of T1-weighted structural images to the mean fMRI image, using normalized mutual information as a cost function. The co-registered anatomical images were segmented into gray matter, white matter, cerebro-spinal fluid and non-brain tissue using New Segment and nonlinearly registered to MNI152 space. Normalized functional images underwent a Gaussian smoothing with a full-width-at-half-maximum of FWHM=8mm.

#### c) Resting-state fMRI

We employed preprocessing methods described in prior work (Paquola, Seidlitz et al. 2020, Royer, Paquola et al. 2020), and preprocessed the resting-state fMRI using a combination of AFNI (Cox 1996) and FSL (Jenkinson, Beckmann et al. 2012). In brief, the first five TRs were discarded to allow for signal equilibrium and P-A/A-P blipped scan pairs were used to correct for geometric distortions. Following motion correction, we warped the corrected images to the T1-weighted space using a combination of rigid body and boundary-based registrations (Greve and Fischl 2009). A high-pass filter was used to correct the time series for scanner drifts. Additional noise components were removed using ICA-FIX, trained on in-house data from the same scanner (Salimi-Khorshidi, Douaud et al. 2014). The resting-state time series were sampled at each cortical vertex and along hippocampal subfields surfaces, as described previously (Bernhardt, Bernasconi et al. 2016). The cortical time series were mapped to the native surface and subsequently registered to Conte69 template using tools from connectome workbench.

### Gradient identification

We followed previous approaches to identify neocortical (Margulies, Ghosh et al. 2016) and hippocampal connectivity gradients (Vos de Wael, Lariviere et al. 2018). Briefly, we computed the Pearson correlation coefficient map between timeseries of all pairs of cortical vertices for each subject. Correlations underwent Fisher r-to-z transformations, and maps were averaged across subjects. The group mean functional connectivity matrix was converted back to *r* values using a hyperbolic tangent function that scales them between −1 to 1. The top 10% connections for each row were retained and all others were set to zero. A cosine similarity matrix was calculated followed by diffusion map embedding, a nonlinear dimensionality reduction technique, to identify principal modes of cortical spatial variations in functional connectivity (Coifman and Lafon 2006). Hippocampal gradient computations were similar to the computations for neocortical gradients, except that connectivity patterns were calculated from hippocampal vertices to cortical parcels (Glasser, Coalson et al. 2016), and that a normalized angle affinity matrix was computed instead of a cosine similarity matrix as in previous work (Vos de Wael, Lariviere et al. 2018). Neocortical and hippocampal gradients were aligned to normative gradients obtained from the Human Connectome Project dataset using Procrustes rotation (Langs, Golland et al. 2015). All gradient computations were performed using the BrainSpace toolbox, which is openly available at https://github.com/MICA-MNI/BrainSpace (Vos de Wael, Benkarim et al. 2020).

### Pattern separation analysis in healthy individuals

#### a) Functional activation mapping

We employed an event-related fMRI analysis, and convolved trial-specific delta functions with the canonical hemodynamic response function (the sum of two gamma functions). For each participant, each of the two event types, (*similar*|*lure*) and (*similar*|*novel*) conditions, was modelled by a regressor of interest, and 6 motion parameters were included as nuisance regressors in a voxel-wised general linear model. Contrasts were created for each subject to address pattern separation, defined by (*similar*|*lure*) - (*similar*|*novel*) One sample t-tests examined effects at the second (*i.e*., group-wise) level. Contrasts for the two conditions at the first (*i.e*., subject-specific) level analysis, and the *t-*value map for the one-sample *t* test describing group effects, were mapped onto cortical and hippocampal surfaces using a boundary-based registration procedure (Greve and Fischl 2009).

#### b) Gradient profiling

We stratified findings based on the previously mapped connectivity gradients in hippocampal (Vos de Wael, Lariviere et al. 2018) and neocortical regions (Margulies, Ghosh et al. 2016) to associate the spatial distribution of effects with respect to main axes of functional organization. In brief, we computed Spearman correlations between pattern separation activation t-values and first/second gradients for the neocortex and for hippocampal subfields respectively. To account for spatial autocorrelations, we used non-parametric spin tests (Alexander-Bloch, Shou et al. 2018) implemented in BrainSpace (Vos de Wael, Benkarim et al. 2020). Only positive t-values were considered.

#### c) Single subject analysis

In addition to group-wise analysis, we verified results at the single subject level. To this end, we mapped subject-specific *t*-maps to the gradient space, thresholded these (t>0), and assessed the correlation between individual activation patterns and neocortical as well as hippocampal gradients.

### Comparison between healthy controls and patients

Patients underwent the same task-based and resting-state fMRI scans as healthy controls.

#### a) Behavioral comparisons

We compared BPS scores between patients and controls, adjusting for age and sex. We also compared each of the nine different item/response combinations (*i.e*., repetition items labeled as old/similar/new, lure items labelled as old/similar/new, novel items labelled as old/similar/new). Findings were corrected for multiple comparisons using the false discovery rate (FDR) procedure with *q* = 0.05 (Benjamini and Hochberg 1995).

#### b) Comparison of functional activation for pattern separation

For each group, we had the contrast map representing the differences between betas of conditions (*similar*|*lure*) and (*similar*|*novel*) mapped on neocortical and hippocampal surfaces, controlling for age and sex.

For each group, we computed spearman correlations between task-related *t*-values and functional gradients across neocortical and hippocampal vertices in both the left and right hemisphere separately. As determined above, significances were determined using non-parametric spin tests that control for spatial autocorrelation (Alexander-Bloch, Shou et al. 2018). Between group comparison in correlations were performed using spin permutation test to compare the real group difference with null distribution of the group difference.

#### c) Associations to BPS

Neocortical and hippocampal *t*-value maps for all subjects were weighted by the corresponding gradient to generate a single scalar loading score. This provided a personalized loading score, with high loadings indicating that activity patterns followed the principal gradients, and low loadings the opposite. We then calculated regressions between individual differences in residual BPS scores and neocortical as well as hippocampal loadings, respectively.

## Acknowledgements

Qiongling Li was funded by China Scholarship Council Scholarship. Casey Paquola was funded through a postdoctoral fellowship of the Fonds de la Recherche due Quebec – Santé (FRQS). Bo-yong Park was funded by Molson Neuro-Engineering fellowship by Montreal Neurological Institute and Hospital (MNI). Jessica Royer was funded by a fellowship from the Canadian Institutes of Health Research (CIHR). Reinder Vos de Wael was supported by the Savoy Foundation for Epilepsy Research. Andrea Bernasconi and Neda Bernasconi were supported by FRQ-S and CIHR (MOP-57840, MOP-123520). Birgit Frauscher receives funding from FRQ-S (Chercheur-Boursier clinician Junior 2) and the National Science and Engineering Research Council of Canada (NSERC Discovery and Accelerator Supplement). Jonathan Smallwood was funded the European Research Council (WANDERINGMINDS). Lorenzo Caciagli acknowledges support from the NINDS (R01-NS099348-01) and from a Berkeley Fellowship (UCL and Gonville and Caius College, Cambridge). Boris Bernhardt acknowledges research support from the NSERC (Discovery-1304413), the Canadian Institutes of Health Research (CIHR FDN-154298), SickKids Foundation (NI17-039), Azrieli Center for Autism Research (ACAR-TACC), and the Tier-2 Canada Research Chairs program.

